# Biocide’s susceptibility and frequency of biocide resistance genes and the effect of exposure to sub-inhibitory concentrations of sodium hypochlorite on antibiotic susceptibility of *Stenotrophomonas maltophilia*

**DOI:** 10.1101/2021.10.04.463145

**Authors:** Raana KazemzadehAnari, Amir Javadi, Farhad Nikkhahi, Mehdi Bakht, Mohammad Rostamani, Akram Azimi, Fatemeh Zeynali Kelishomi, Safar Ali Alizadeh

**Affiliations:** Medical Microbiology Research Center, School of Medicine, Qazvin University of Medical Sciences, Qazvin, Iran; Department of Community Medicine, School of Medicine, Qazvin University of Medical Sciences, Qazvin, Iran

**Keywords:** *Stenotrophomonas maltophilia*, Biocides, Antibiotic resistance, nosocomial infection, sub-MIC, QACs, efflux pump

## Abstract

The overuse of biocides in healthcare-facilities poses risk for emergence and spread of antibiotic resistance among nosocomial pathogens. Hospital-acquired infections due to *S. maltophilia*have been increased. The objective of this study was to evaluate the susceptibility of *S. maltophilia* clinical isolates to commonly used biocides in hospitals, as well asthe frequency of biocides resistance genes among them. This study also intended to assess the effect of exposureof *S. maltophilia* isolates to sub-inhibitory concentrations of sodium hypochlorite upon the antimicrobial susceptibility patterns.This study included 97 *S.maltophilia*isolates.Biofilm formation was determined by microtiter plate assay. The susceptibility tests of five biocides were studied against all *S. maltophilia* isolates by microbroth dilution method. Susceptibility of isolates to antibiotics by disk diffusion method were compared before and after exposure to sub-inhibitory concentrations of sodium hypochlorite. Presence of *qacE, qacE*Δ*1, SugE* genes was screened by PCR. Based on minimum inhibitory and bactericidal concentrations of biocidessodium hypochlorite 5% and ethyl alcohol 70% were the strongest and weakest against *S. maltophilia* isolates, respectively. The frequency of *sugE* gene resistance genes was found 90.7% in our clinical *S. maltophilia* isolates. None of the isolates carried *qacE* and *qacEΔ1* gene. Exposure to sub-inhibitory concentration of sodium hypochlorite showed significantly change the susceptibility of isolates towards ceftazidime (*P* = .019), ticarcillin/clavulanate (*P* = .009), and chloramphenicol (*P* = .028).This study demonstrated that exposure to sub-inhibitory concentrations of sodium hypochlorite leads to reduced antibiotic susceptibility and development of multidrug-resistant *S.maltophilia* strains.

## Introduction

Hospital-acquired infections are recognized as one of the problematic challenges for infection control worldwide.Healthcare facilities and environments provide an idealreservoir for the growth, colonization, and proliferation of pathogenic organisms(1, 2).*Stenotrophomonas maltophilia*, formerly known as *Pseudomonas maltophilia* or *Xanthomonas maltophilia*is a common cause of hospital-acquired infection(3).Despitelimited pathogenicity and due to intrinsic resistance nature and acquiring resistance of this bacterium against multiple antimicrobial agents and biocides throughplasmids, transposons, integrons, and limited therapeutic options,*S. maltophilia* is known as one of the leading antibiotic-resistant pathogens and isassociated with avariety of life-threatening nosocomial infections in hospitalized or immunocompromised patients(3, 4).Additionally, the ability to adhere and develop biofilm both on biotic and abiotic surfaces and survive in adverse environmental conditions, enables *S. maltophilia* to causes infection and contributes to chronic infections. Resistance to antimicrobial agents is the most important property of biofilm development(5, 6).

The increasing prevalence of biocide resistance and the potential for cross-resistance to some antibiotics is one of the global health threats and result in hospital-acquired infections and ineffective treatments. The development of resistance to biocides by bacteria could be a public health hazard. Biocides including antiseptics and disinfectantswith proper use, are an essential part of public health and have a crucial role in preventing colonization and infection and controlling pathogenic bacteria in thehospital. The effectiveness of biocides depends on several factors such as concentration, thestatus of bacteria (biofilm or planktonic), and presence of genes conferring resistance to biocides(7). Based on mentioned points, there is a rational concern that the misuse of biocides such as highor inadequate concentrations, and frequent exposure to sub-inhibitory concentrations (concentrations below those required to arrest growth) could select for strains that are tolerant to and could render them ineffective and may contribute to antibiotic resistance and leads to the development of multi-drug resistant (MDR) strains(8). One of the well-known mechanisms responsible for resistance to biocides is the expression of efflux systems involving *qac* genes(*qacE, qacE*Δ*1*) and also *sugE* gene(9). These resistance genes are members of the SMR family which conferring resistance to QACs(9–11). The co-resistance and cross-resistance to biocides and antibiotics could be relevant to genes encoding resistance to biocide horizontally transferred mobile genetic elements that also carry antibiotic resistance genes(12). Since as, few published studies are available to assessing reduced susceptibility to biocides than antibiotics and also about antibiotic resistance induced by increased resistance to biocides against *S. maltophilia*.We aimed to investigate the susceptibility of *S. maltophilia* isolates to biocidesas well as the prevalence of biocides resistance genes andthe effect of exposure to sub-inhibitory concentrations of the sodium hypochlorite on antimicrobial susceptibility patterns of *S. maltophilia* clinical isolates in Iran. Undoubtedly, theresults of this study and understanding thesusceptibility of *S. maltophilia* to biocides andits correlation with antibiotic resistance will contribute to the control of this bacterium in hospitals and aid in the prevention of nosocomial infection.

## Material and method

### Isolation and identification

A total of 105 *S. maltophilia* clinical isolates were collected in the present study during the period between September 2019 and March 2020 at five tertiary-care hospitals (H1-H5) in Iran(Tehran and Qazvin).All of the isolates were identified using standard microbiological and biochemical methods such as Gram stain, catalase and oxidase tests, motility, oxidative or fermentative metabolism, deoxyribonuclease test agar (DNase), triple sugar iron agar (TSI), lysine decarboxylase and esculin hydrolysis (Merck, Germany)(13). Genomic DNA was extracted from asingle colony of each isolate with high pure PCR Template Preparation Kit (Roche company, Germany). All isolates were reconfirmed genotypically as *S. maltophilia* by PCR with specific primers illustrated in Table 1 to amplify a 638-bp fragment of the *23S rRNA* gene and stored at −20°C in trypticase soy broth (TSB; Merck, Germany) supplemented with 20% glycerol for further analysis. *Pseudomonas aeruginosa* ATCC 27853 and *S. maltophilia* ATCC 13637 were used as the quality control strains. A representative amplicon of *23S rRNA* gene was subjected to sequencing andthe sequence was deposited in GenBank and assigned the accession no**<u>MZ468054</u>**.

**TABLE 1.**
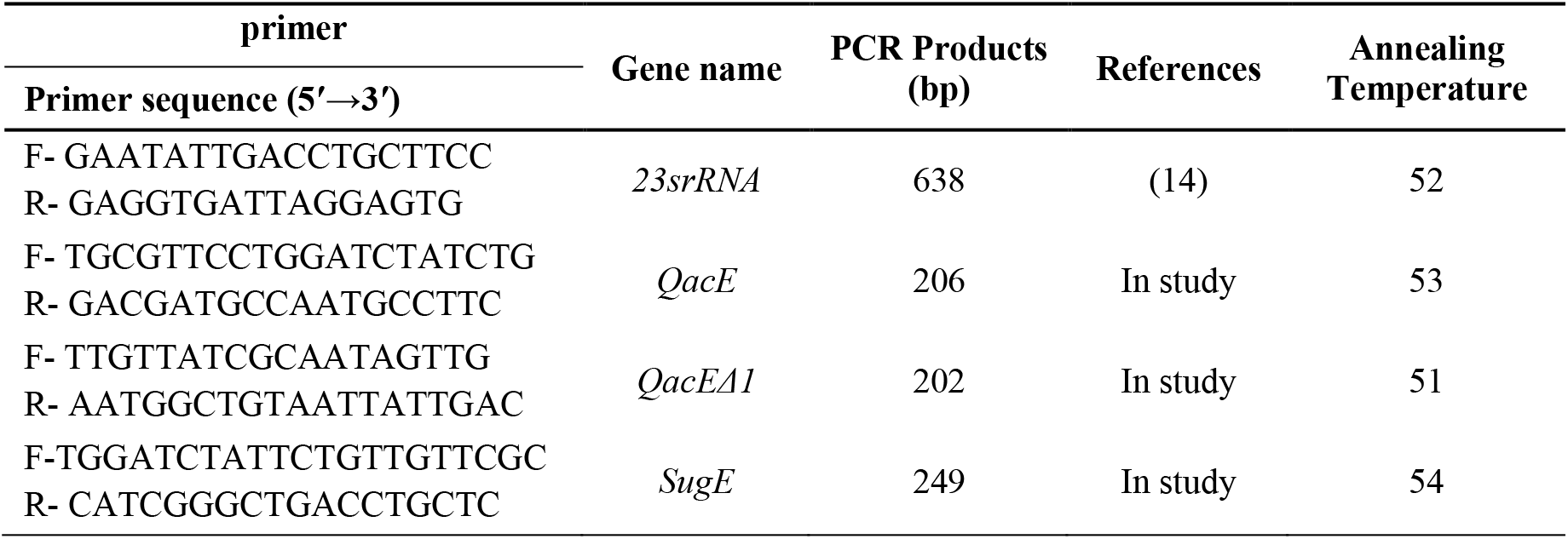
List of primers used in the study

### Antibiotic susceptibility testing

Antibiotic susceptibility testing of *S. maltophilia* isolates against meropenem (10 mg), imipenem (10 mg), trimethoprim/sulfamethoxazole (1.25/23.75mg), levofloxacin (5 mg), and minocycline (30 mg) (Mast Group Ltd, UK) was determined using Kirby-Bauer disc diffusion method according to the criteria of the Clinical and Laboratory Standards Institute (CLSI) guidelines(15). The critical breakpoints of ceftazidime (30 mg) and ticarcillin/clavulanate (75/10 mg) of *Pseudomonas aeruginosa* were used for interpretation of the results because no breakpoints for *S. maltophilia* were recommended by the CLSI. The results of chloramphenicol (30 mg) and tigecycline (15 mg) were interpreted according to the CLSI breakpoints of Enterobacteriaceae and the Food and Drug Administration (FDA), respectively. The *P. aeruginosa* ATCC 27853 and *S. maltophilia* ATCC 13637 were used for susceptibility testing. Due to theintrinsic resistance nature of *S. maltophilia*to carbapenems, susceptibility to meropenem and imipenem was also determined to confirm the identity of the isolates.*S. maltophilia* isolate was defined as multidrug-resistant (MDR), if it exhibited non-susceptibility to at least one agent in three or more antimicrobial categories including β-lactam/β-lactamase inhibitor combinations, sulfonamides, fluoroquinolones, chloramphenicol, cephalosporins, tetracyclines, and glycylcyclines(16).

### Biocides tested

During this study five commonly used antiseptics and disinfectants in hospitalsfor clinical items and bio-cleaning of instruments and surfaces were subjected to testingincluding: Ethyl Alcohol (96% v/v, JATA Co., Arak, Iran) was used by diluting the stock solution with sterile distilled water to 70% (v/v), Dettol Antiseptic Disinfectant Liquid (Chloroxylenol 4.8% w/v, Reckitt and Benckiser Ltd. India), Sodium hypochlorite (containing 5% w/v active chlorine, Golrang co., Iran), Chlorhexidine (2% w/v, Maquira co; Brazil), Sayasept-HP 2% (Fifth-generate QACs, didecyl dimethyl ammonium chloride (DDAC) and alkyl dimethyl benzyl ammonium, Bs co., Iran).

### Determination of minimum inhibitory and bactericidal concentrations (MICs/MBCs) of biocides

Susceptibility testing for biocides was performed using a modified broth microdilution method(17). In brief, in the beginning, the *S. maltophilia*isolates were grown overnight on Muller Hinton agar(Merck, Germany) at 37°C*.S. maltophilia* suspensions were adjusted to a turbidity equivalent to that of a 0.5 McFarland standard with sterilized saline solution and then diluted 0.01%. The wells 1 to 9 of a sterile 96-well plate were filled with 100 μl of trypticase soy broth (TSB). To well 1, 100 μl of tested biocides were added, upon mixing well, two-fold Serial dilutions of biocides were done in TSB to yield the desired concentration ranging from 2 to 512 μg/mL, followed by 100 μl of each tested isolate (1.5 × 10^6^ CFU/mL) were inoculated to wells 1 through 9 to each well. The wells 10 and 11 were growth (TSB + inoculation) and sterility (contained TSB alone) controls, respectively. The final concentration of each well was equal to 5× 10^5^ CFU/mL. MICs were examined visually after incubation at 37 °C for 24 h. The lowest concentration of the tested biocides that inhibited visible bacterial growth and didn’t show turbidity was reported as the minimum inhibitory concentration (MIC).To determine the minimum bactericidal concentration (MBC), approximately 100 μL was withdrawn from each well without visible bacterial growth were cultured onto Muller Hinton agar plates and incubated overnight at 37°C.The MIC and MBC of each biocide for all 97 strains of S. *maltophilia* were determined using this method.

### Effect of exposure to sodium hypochlorite on antibiotic susceptibility of the isolates

The effect of sub-inhibitoryconcentrations of used sodium hypochloritefollowing exposure on the susceptibility of *S. maltophilia* isolates was evaluated by comparing the antibiotics susceptibility patterns of isolates before and after exposure toasub-inhibitory concentration of sodium hypochlorite. Briefly, after determining the MIC and MBC, bacterial suspensions were withdrawn from wells containing the highest concentration of sodium hypochlorite which still allows bacteria to grow (sub-MIC), and were cultured onto Muller-Hinton agar plates and incubated overnight at 37°C to isolate the survived organisms. The antibiotic susceptibility test of those survived *S. maltophilia* isolates was performed again by disk diffusion method following exposure to the sub-inhibitory concentrations of sodium hypochlorite. The results were compared with previous results.

### Biofilm formation assay

Biofilm assay was conducted in triplicate in 96-well flatbottomed polystyrene microplates to evaluatethe capacity of biofilm production in *S.maltophilia* isolates as described previously with some modifications(18). Initially, the bacterial suspensions were prepared with an optical density (OD)of 0.1 were adjusted by using sterile trypticase soy broth (TSB) at 600 nm (OD600) with a spectrophotometer. Then, 200 μl of adjusted inoculums were transferred in triplicate into sterile 96-well flat-bottomed microplates and incubated overnight in a 37°C. A series of triplet wells contained TSB alone (uninoculated broth) was considered as negative. The media were then removed by slightly tapping the plate and washed three times with phosphate-buffered saline (PBS: pH 7.2).Adherent biofilms were fixed with methanol for 15 min and dried at room temperature. Then, the biofilms within the wells were stained with 200 μl of theaqueous solution of 1% (w/v) crystal violet for 15 min. To remove the dye attached to the biofilm layers, the wells were rinsed three times with PBS and the plate was air-dried, biofilms were detached by adding 200 μl of 33% acetic acid into each well for 15 min. The optical density was measured at 570 nm (OD570) using a microtiter plate reader (BioTek, Epoch, USA). The point to be noted isthat all experiments were carried out in triplicate and repeated three times. Additionally, the optical densitycut-off value (ODC) was established and defined as three standard deviations (S.D) above the mean OD of the negative control. (ODC = the average OD of the negative control + 3 × S.D of the negative control). The isolates were classified into four groups based upon the strength of biofilm formation as follows(19): no biofilm production (OD ≤ ODC); weak biofilm-producer (ODC <OD ≤ 2 × ODC); moderate biofilm-producer (2×ODC < OD ≤ 4×ODC); and strong biofilm-producer (ODC <4OD).

### Screening and detectionof*qacE*, *qacEΔ1* and *sugE*

The presence of *qacE*, *qacEΔ1*, and *sugE*genes that confer resistance to biocides was examined using the primers shown in Table1. PCRs were conducted on a thermal cycler (Applied Biosystems, USA) in 25 μl reaction volume containing 10 μL of 2X Master Mix RED (Ampliqon, Denmark), 1 μl of 10 pmol of each primer (Sinaclon Co; Tehran, Iran), 50 ng of template DNA and 6 μl of sterile distilled water. PCR conditions were performed under the following thermal conditions: pre-denaturation at 94 °C for 5 min; 30 cycles of DNA denaturation for 1 min at 94 °C; annealing at 51-54°C, according to the primers for each gene (SupplementaryTable 1) for 25 s, extension for 50 s at 72°C and a final extension at 72°C for 7 min. All of the amplified products were separated by electrophoresis in 1.8% agarose gel with 1× TBE bufferand visualized under ultraviolet light. A representative PCR amplicon of each gene was sequenced to ensure the specific amplification and analyzed using BLAST (http://www.ncbi.nlm.nih.gov/BLAST/).

### Statistical analysis

Data are expressed frequency and percent. Pearson chi-square or Fisher’s exact test was used to determine significant differences between proportions. The non-parametric Wilcoxon signed-rank test was performed to comparison of the antibiotics’ patterns before and after exposure of *S. maltophilia* isolates to sodium hypochlorite.The values P < 0.05 were considered statistically significant.Statistical analysis was done using SPSS version 16.0 statistical software (SPSS Inc., Chicago, IL, USA).

### Data availability

A representative amplicon of the *sugE* gene was submitted to the GenBank database under accession number <u>**MZ503513**</u>.

## Result

Biochemical tests and thepresence of a 638-*bp* fragment of *23S rRNA gene*in 97 test isolates confirmed their identity as *S. maltophilia* and none of the isolates were removed from the study. Out of 97 isolates, 55 (56.7%) and 7 (7.2%) isolates were collected from hospitals H1, H2 affiliated to Tehran University of Medical Sciences (TUMS) and 25 (25.8%) isolates from H3, 8 (8.2%) from H4 and H5 with 2 (2.1%) isolates from admitted patients in hospitals affiliated to Qazvin Medical University (QUMS). Among them, 59 (60.8%) isolates were from males and 38 (39.2 %) isolates were from females (male: female ratio =1.5). The range of patients’ age was from 2 days to 85 years and 9 (9.3%) of the isolates were recovered from infants and 3 cases of whom were from infants less than one month of age (<1).Blood was the major source of isolates (n =82; 84.5%) and the remaining isolates were recovered from tracheal aspirate (n=6;6.2 %) followed by bronchoalveolar lavage (n = 3; 3.1 %), ocular discharge (n= 2; 2.1%), sputum (n=2; 2.1%), urine (n=1; 1%), and ascites fluid (n=1; 1%) (Table 4). The majority of *S.maltophilia* isolates (n = 57; 58.8%) were obtained from patients admitted to emergency wards.

### Antibiotic resistance phenotypes

The antibiotic susceptibility pattern, using the disk diffusion method, of the 97 *S. maltophilia* isolates before treatment with biocides is shown in Table 3. Of the 97 *S. maltophilia* isolates, 25 (25.8%) were multidrug-resistant and 10 (10.3%) isolates were extensively drug-resistant according to CLSI interpretive criteria. As shown in Table 3, among *S. maltophilia* isolates examined, all of them were highly resistant (100%) to imipenem and meropenem, and 10 (10.3%) isolates showed resistance to trimethoprim/ sulfamethoxazole (TMP–SXT) and 4 (4.1%) indicated intermediate resistance. Levofloxacin, minocycline, and tigecycline exhibited the highest susceptibility of 97.9, 88.7, and 86.6%, respectively. The point to be mentioned is that in this studyintermediate susceptibility isolates were intended to be resistant. The resistance rates of isolates to other antibiotics by disk diffusion were as follows: ceftazidime (76.3%); ticarcillin/clavulanate (60.9%); chloramphenicol (39.2%).

### Biocide susceptibility testing

The susceptibility of five biocides was tested against 97 *S. maltophilia* isolates using concentrations ranging from 2 to 512 μg/ml (50%–0.19%). The obtained MIC and MBC results for all of the isolates are shown in Table 2. All of the tested biocides except ethyl alcohol, at MIC and MBC 2-8 had acompleteinhibitory and lethal effect on the *S. maltophilia* isolates whereas, ethyl alcohol at MIC 8 had failed to impede the growth of the isolates. As table shows, the MIC values of the biocides tested were quite variable and in the following ranges: from 64 to 512 μg/mL for sodium hypochlorite, 64 to 256 μg/mL for Dettol, 32 to 256 μg/mL for chlorhexidine, 16 to 128 μg/mL for sayasept-HP and 8 to 128 μg/mL for ethyl alcohol. The isolates with sodium hypochlorite and Dettol MICs of 128 μg/ml, sayasept-HP and chlorhexidine MICs of 64 μg/ml, and ethyl alcohol MICs of 32 μg/ml were observed often.

**TABLE 2.** Summary of biocides MIC and MBCs values determined for 97 *S. maltophilia* isolates

| Biocides | Starting concentrations | Dilution rang tested |  |  |  |  |  |  |
| --- | --- | --- | --- | --- | --- | --- | --- | --- |
|  |  | 1/8 | 1/16 | 1/32 | 1/64 | 1/128 | 1/256 | 1/512 |
| MIC |  |  |  |  |  |  |  |  |
| Ethyl Alcohol | 70% | 2 (2.1%) | 38 (39.2%) | 41 (42.3%) | 15 (15.5%) | 1 (1%) | 0 | 0 |
| Saya sept-HP | 2% | 0 | 2 (2.1%) | 38 (39.2%) | 41(42.3%) | 16 (16.5%) | 0 | 0 |
| Chlorhexidine | 2% | 0 | 0 | 17(17.5%) | 42 (43.3%) | 37 (38.1%) | 1 (1%) | 0 |
| Dettol | 4.8% | 0 | 0 | 0 | 32 (33.0%) | 50 (51.5%) | 15 (15.5%) | 0 |
| Sodium hypochlorite | 5% | 0 | 0 | 0 | 24 (24.7%) | 43 (44.3%) | 27(27.8%) | 3 (3.1%) |
| MBC |  |  |  |  |  |  |  |  |
| Ethyl Alcohol | 70% | 35 (36.1%) | 32 (33%) | 30 (30.9%) | 0 | 0 | 0 | 0 |
| Sayasept- HP | 2% | 0 | 21 (21.6%) | 49 (50.5%) | 26 (26.8%) | 1 (1%) | 0 | 0 |
| Chlorhexidine | 2% | 0 | 0 | 44 (45.4%) | 41 (42.3%) | 11 (11.3%) | 1 (1%) | 0 |
| Dettol | 4.8% | 0 | 0 | 11 (11.3%) | 56 (57.7%) | 28 (28.9%) | 2 (2.1%) | 0 |
| Sodium hypochlorite | 5% | 0 | 0 | 0 | 45 (46.4%) | 36 (37.1%) | 15(15.5%) | 1 (1%) |

The MBCs ranged from 64- 512 μg/mL for sodium hypochlorite, 32-256 μg/mL for Dettol and chlorhexidine, 16-128 μg/mL for sayasept-HP and 8-32 μg/mL for ethyl alcohol in *S. maltophilia* isolates. The isolates with sodium hypochlorite and Dettol MIBs of 64 μg/ml, sayasept-HP and chlorhexidine MBCs of 32 μg/ml, and Ethyl Alcohol MICs of 16 μg/ml were observed most frequently.

### Effect of exposure to sub-inhibitory concentrations of Sodium hypochlorite on antibiotic susceptibility pattern

The antibiotic susceptibility of isolates was retested using disk diffusion only for antibiotic susceptible and susceptibility-intermediate isolates following exposure to sub-inhibitory concentrations of sodium hypochlorite. The obtained results before exposure to biocide were compared to those after exposure. The susceptibility patterns of some isolates either changed from susceptible to susceptibility-intermediate and resistant and from susceptibility-intermediate to resistant. Exposure to thesub-inhibitory concentration of sodium hypochlorite showed asignificant changein the susceptibility of isolates towards ceftazidime (*P* = .019), ticarcillin/clavulanate (P = .009), and chloramphenicol (P = .028), which was susceptible or susceptibility-intermediate to them before exposure, whereas isolates did not show any difference in the susceptibility patterns of the other antibiotics upon exposure to sub-MICs of sodium hypochlorite. Exposure to thesub-inhibitory concentration of sodium hypochlorite showed asignificant increase (P= .014) in the frequency of MDR and XDR *S. maltophilia* isolates. Thus,29 (29.9%) were multidrug-resistant and 11 (11.3%) isolates were extensively drug-resistant. Table3summarizes the susceptibility results tested before and after exposure with sub-MICs of sodium hypochlorite.

**TABLE 3.**
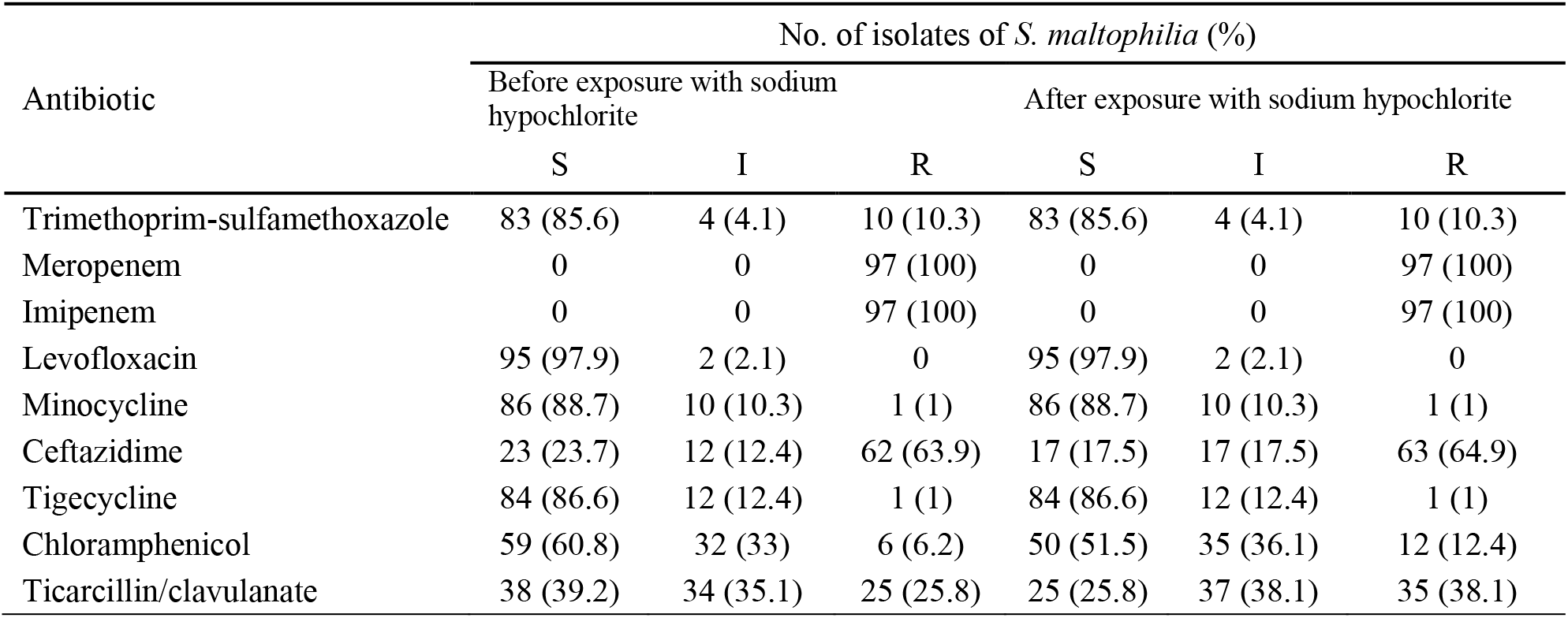
Antibiotic susceptibility profiles of S. *maltophilia*before and after exposure to sub-inhibitory concentrations of sodium hypochlorite

### Biofilm formation

The ability to develop biofilm varied greatly among the *S. maltophilia* isolates. Biofilm phenotypes accounted for 93 out of 97 isolates (95.9 %). The results of the biofilm formation assay showed that 45 (46.4%) of isolates were strong biofilm producers, 41 isolates (42.3.%) were moderate biofilm producers and 7 isolates (7.2%) were weak biofilm-producers; whereas, 4 isolates (4.1%) did not form biofilm. Also, in thepresent study statistical analysis to evaluate the association between antibiotic resistance and biofilm production showed that between antibiotic resistance of ceftazidime (P = 0.049), ticarcillin-clavulanic acid (P = 0.00), and biofilm production was found to be statistically significant. This finding was not seen with other antibiotics.

**TABLE 4.** Occurrence of *S. maltophilia* biofilm in relation to clinical source

| Clinical source (no. of isolates) | No. (%) of isolates with biofilm phenotype |  |  |  |
| --- | --- | --- | --- | --- |
|  | None | Weak | Intermediate | strong |
| blood culture (82) | 4 (4.1) | 3 (3.09) | 39 (40.2) | 36 (37.1) |
| urine culture (1) | - | - | - | 1 (1.03) |
| Trachea culture (6) | - | 2 (2.06) | 1 (1.03) | 3 (3.09) |
| BAL (3) | - | 1 (1.03) | - | 2 (2.06) |
| Eye discharge culture (2) | - | - | - | 2 (2.06) |
| Sputum (2) | - | 1 (1.03) | 1 (1.03) | - |
| Ascites (1) | - | - | - | 1 (1.03) |
| Total (97) | 4(4.1) | 7 (7.2) | 41 (42.3) | 45 (46.4) |

BAL, bronchoalveolar lavage

### Distribution of Biocides Resistance Genes

PCR screening showed that among the 97 isolates tested, *sugE* gene that confers resistance to biocides was present in 88 (90.7%) isolates. Whereas, the *qacE* and *qacEΔ1* genes were not detected in any of the isolates.

**Fig1:**
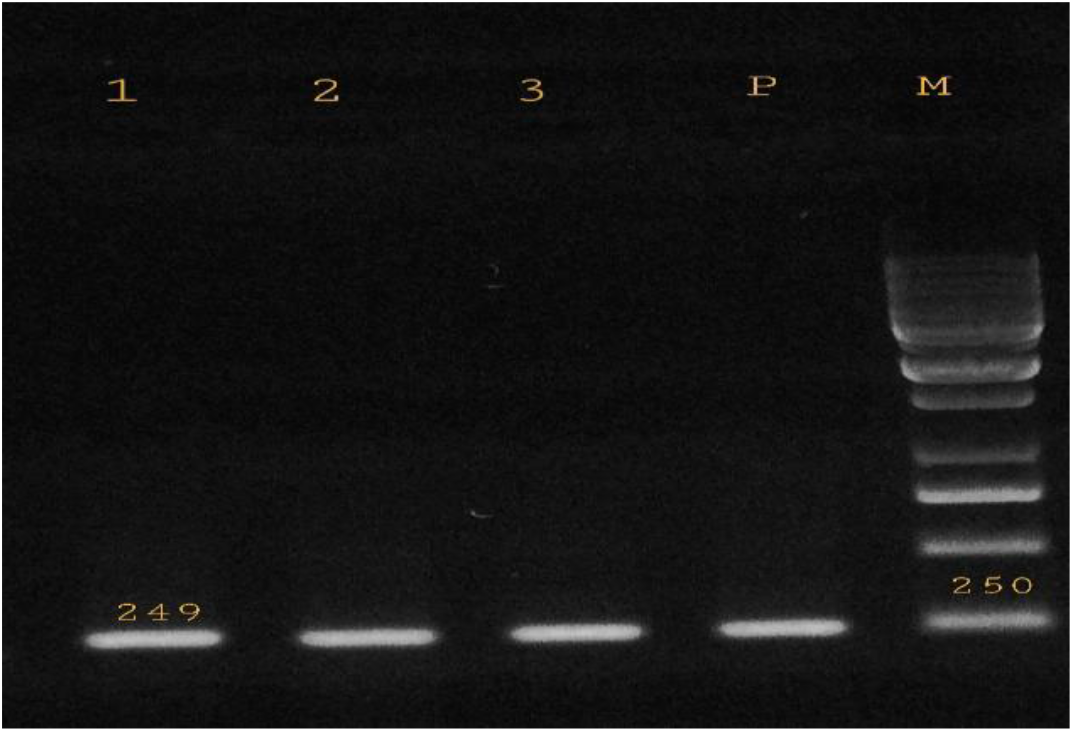
Gel electrophoresis of the PCR amplified products of *sugE* gene for the *S. maltophilia* isolateswith 249 bp amplification fragment. Lane M: DNA size marker - Lane P: positive control - Lane 1-3: *sugE* positive isolates.

## Discussion

Biocides with proper use, have a crucial role in preventing colonization and infection of pathogenic microorganisms. The overuse and suboptimal concentrations of biocides used in hospitals for infection control might contribute to increased MICs of them and leads tothe development of resistance to biocides and multi-drug resistant strains(8). Cross-resistance between antibiotics and biocides may occur via various and common mechanisms between them such as efflux pump systems, changes in permeability, and biofilm formation(20).

Since as, the increasing emergence of bacteria with reduced susceptibility to biocides and the possible linkage between biocide and antibiotic resistance is a newly important concern, in this study we attempted to find a possible association between reduced susceptibility to biocides and the presence of biocide resistance genes in clinical isolates of *S. maltophilia*. Here, the susceptibility to biocides was determined by comparing the MIC and MBC values of a set of 97 isolates of *S. maltophilia* using broth microdilution and designed a 96 well plate to accommodate five biocides containing nine two-fold dilutions of each biocide. Since there are no established breakpoints available for MICs and MBCs for defined resistance to biocidesagainst *S. maltophilia*, we tested a 2-Fold dilution from 50% to 0.19% of each biocide. In the present study, the biocides selected for susceptibility testing were sodium hypochlorite 5%, Dettol 4.8%,chlorhexidine2%, sayasept-HP 2%, ethyl alcohol 70%, because they are widely used as antiseptic and disinfectants in healthcare facilities in Iran. Given that, there aren’t any criteria for categorization of bacteria as susceptible or resistant to biocides according to MIC and MBC data, we found that among tested biocides, sodium hypochlorite 5% (the lowest MIC and MBC) and ethyl alcohol 70% (the highest MIC and MBC) were the strongest and weakest against *S. maltophilia* isolates, respectively. Overall, based on theminimum inhibitory and bactericidal concentration values, the most effective biocides were sodium hypochlorite 5%, Dettol 4.8%,chlorhexidine 2%,saya sept-HP 2%, ethyl alcohol 70%, respectively.

The *sugE* gene along with*qacE/EΔ1* genes which are members of small multidrug resistance (SMR) protein is also being known as a quaternary ammonium compound (QAC) resistance determinant. As far as we know, a limited number of biocides resistance gene studies have involved clinical *S. maltophilia* isolates. In this study, the frequency of *qacE*, *qacΔE1*, and *sugE* genes was assessed and this is the first study in Iran that investigated thesusceptibility of biocides against *S. maltophilia* isolates and showing thedistribution of biocides resistance genes among them.C. wang et al found that 2 out of 19 (10.5%) *S. maltophilia* isolates carried *qacΔE1*gene and inthe study of Huang (21) the prevalence of *qacEΔ1*gene was reported 10.5%. Kückenet alfound any *qacEΔ1* and *qacE* gene out of 13 *S. maltophilia* isolate(9). However, in our study *qacE* and *qacEΔ1* genes were not detected in any isolates.In contrast, our findings have demonstrated high levels of presence of *sugE* gene (90%) in clinical isolates of *S. maltophilia* whilethere was no significant correlation between thepresence or absence of *sugE* gene and increased MICs and MBCs of tested biocides against *S. maltophilia isolates.*

Biofilms are associated with 65% of hospital-acquired infections(22). Reports suggesting that biofilm formation is an important mechanism for resistance to antibiotics and biocides by*S. maltophilia*(23).Here, we observed that all but four isolates investigated were biofilm-producers, although with different biofilm-forming abilities. The prevalence of S*. maltophilia* isolates able to develop biofilm in our study (95.9%) was like that (88.7–100%) in previous reports in Iran and Europe(19, 22, 24, 25). Also, the present study examined the association between antibiotic resistance andpotential of biofilm formation, these results demonstrated that there is a significant association between the potential of biofilm formation and resistance to ceftazidime and ticarcillin/clavulanate in *S. maltophilia*, whichwas consistent with the report by Sun et al(26).

Some previous studies have demonstrated that antibiotic resistance can be induced by sub-MICsconcentrations of biocide(27). The effect of biocides on antibiotic susceptibility in bacteria and thedevelopment of antibiotics-resistanthealthcare-acquired microorganisms after treatment with sub-MICs of different biocides on surviving bacteria has been measured and confirmed. In this study, we also examined whether sub-MICs of sodium hypochlorite can induce antibiotic resistance in clinical isolates of *S. maltophilia*. In this regard, the antibiotics susceptibility of isolates after exposure to the sub-MICs of sodium hypochlorite was determined and compared with those determined before exposure to sodium hypochlorite. To our knowledge, no previous studies have investigated the effect of exposure to biocides on *S. maltophilia* resistance to antibiotics. Our finding demonstrated that *S. maltophilia* isolates couldyield resistance toward antibiotics after overnight incubation with sodium hypochlorite, a statistically significant change was observed in susceptibility patterns of ceftazidime, ticarcillin/clavulanate, and chloramphenicol. Notably, the number of multidrug-resistant *S. maltophilia* isolates has been shown a statistically significant increase, in comparison to before exposure to the biocide. The results of our study, together with previous studies, suggest that exposure to the sub-inhibitory concentrations of various biocides can induce antibiotic resistance in the isolates.

From the comparison between the obtained results, it can be concluded that bacterial antibiotic resistance is not necessarily a reason for resistance to biocides.In fact, a biocide can have a similar effect on an antibiotic-sensitive or resistant bacteriumand the presence of biocides resistance gene and biofilm are effective in this regard. And also, the present study showed that as long as biocides are used in proper concentrations, they can prevent the growth and development of multi-drug resistant isolates.Whereas using suboptimal concentrations and exposure to sub-inhibitory concentrations of biocides such as sodium hypochlorite result in reduced antibiotic susceptibility and cross-resistance and the development of antibiotic-resistant*S. maltophilia* strains which can cause detrimental effects and increase nosocomial infections.

## Conclusion

In conclusion, our study demonstrated that *sugE* gene was commonly present among clinical *S. maltophilia*. None of the *S. maltophilia* isolates harbored *qacE* and *qacEΔ1*genes. There was no significant correlation between thepresence or absence of *sugE* gene and MICs and MBCs observed in *S. maltophilia* isolates. As far as we know, the present study is the first report showing thedistribution of biocides resistance genes and their biocide MIC and MBC values among *S. maltophilia* isolates from Iran.This study demonstrated that although biofilm-forming capacity was highly conserved among clinical strains of *S. maltophilia*, there wasa significant difference in phenotype among them. This study also showed that exposure tosub-inhibitory concentrationsof sodium hypochlorite leads to reduced antibiotic susceptibility and thedevelopment of multidrug-resistant*S. maltophilia* strains.Consequently, the use of properbactericidal concentrations of different biocides aid in theprevention and controllingthe outbreak of nosocomial infections caused by multi-drug resistant bacteria such as *S. maltophilia.*This study emphasizes the need for using optimal concentrations of biocides and also a large-scale study to evaluate reduced susceptibility to biocides of nosocomial pathogens. The rotational use of different biocides is recommended to avoid the evolution of resistance or selection of resistant strains in the hospital environment.

## Ethical approval

This study was approved by the Ethics Committee of Qazvin University of Medical Sciences “[approval no. IR.QUMS.REC.1398.143]”. Patient information was anonymized.

## Acknowledgments

This research received no specific grant from any funding agency in the public, commercial, or not-for-profit sectors.

## Author contributions

RKA: Designed and performed the microbiologic and molecular experiments, wrote the manuscript. SAA: Supervised the research, corresponding author. FN:The advisor and contributed substantially to the conception and design of experiments. AJ: Methodology and data analysis. MB - MR -AA-FZK: Involved in collecting of samples and performing of experiments. All authors have read and agreed to the published version of the manuscript.

## References

1. Rutala WA, Weber DJ. Disinfection, sterilization, and antisepsis: An overview. American journal of infection control. 2016;44(5):e1–e6.

2. Meade E, Garvey M. Efficacy testing of novel chemical disinfectants on clinically relevant microbial pathogens. American journal of infection control. 2018;46(1):44–9.

3. Brooke JS. Stenotrophomonas maltophilia: an emerging global opportunistic pathogen. Clinical microbiology reviews. 2012;25(1):2–41.

4. Çıkman A, Parlak M, Bayram Y, Güdücüoğlu H, Berktaş M. Antibiotics resistance of Stenotrophomonas maltophilia strains isolated from various clinical specimens. African health sciences. 2016;16(1):149–52.

5. Flores-Treviño S, Bocanegra-Ibarias P, Camacho-Ortiz A, Morfín-Otero R, Salazar-Sesatty HA, Garza-González E. Stenotrophomonas maltophilia biofilm: its role in infectious diseases. Expert review of anti-infective therapy. 2019;17(11):877–93.

6. Pompilio A, Piccolomini R, Picciani C, D’Antonio D, Savini V, Di Bonaventura G. Factors associated with adherence to and biofilm formation on polystyrene by Stenotrophomonas maltophilia: the role of cell surface hydrophobicity and motility. FEMS microbiology letters. 2008;287(1):41–7.

7. Humayoun SB, Hiott LM, Gupta SK, Barrett JB, Woodley TA, Johnston JJ, et al. An assay for determining the susceptibility of Salmonella isolates to commercial and household biocides. PLoS One. 2018;13(12):e0209072.

8. Dynes JJ, Lawrence JR, Korber DR, Swerhone GD, Leppard GG, Hitchcock AP. Morphological and biochemical changes in Pseudomonas fluorescens biofilms induced by sub-inhibitory exposure to antimicrobial agents. Canadian journal of microbiology. 2009;55(2):163–78.

9. Kücken D, Feucht H-H, Kaulfers P-M. Association of qacE and qacE Δ1 with multiple resistance to antibiotics and antiseptics in clinical isolates of Gram-negative bacteria. FEMS Microbiology Letters. 2000;183(1):95–8.

10. Zou L, Meng J, McDermott PF, Wang F, Yang Q, Cao G, et al. Presence of disinfectant resistance genes in Escherichia coli isolated from retail meats in the USA. Journal of Antimicrobial Chemotherapy. 2014;69(10):2644–9.

11. Bay DC, Rommens KL, Turner RJ. Small multidrug resistance proteins: a multidrug transporter family that continues to grow. Biochimica et Biophysica Acta (BBA)-Biomembranes. 2008;1778(9):1814–38.

12. Bennett P. Plasmid encoded antibiotic resistance: acquisition and transfer of antibiotic resistance genes in bacteria. British journal of pharmacology. 2008;153(S1):S347–S57.

13. Adegoke AA, Stenström TA, Okoh AI. Stenotrophomonas maltophilia as an emerging ubiquitous pathogen: looking beyond contemporary antibiotic therapy. Frontiers in microbiology. 2017;8:2276.

14. Whitby PW, Carter KB, Burns JL, Royall JA, LiPuma JJ, Stull TL. Identification and Detection ofStenotrophomonas maltophilia by rRNA-Directed PCR. Journal of Clinical Microbiology. 2000;38(12):4305–9.

15. Wayne P. Clinical and laboratory standards institute. Performance standards for antimicrobial susceptibility testing. 2011.

16. Guan X, He L, Hu B, Hu J, Huang X, Lai G, et al. Laboratory diagnosis, clinical management and infection control of the infections caused by extensively drug-resistant Gram-negative bacilli: a Chinese consensus statement. Clinical Microbiology and Infection. 2016;22:S15–S25.

17. Lotfi M, Vosoughhosseini S, Ranjkesh B, Khani S, Saghiri M, Zand V. Antimicrobial efficacy of nanosilver, sodium hypochlorite and chlorhexidine gluconate against Enterococcus faecalis. African Journal of Biotechnology. 2011;10(35):6799–803.

18. Stepanović S, Vuković D, Hola V, Bonaventura GD, Djukić S, Ćirković I, et al. Quantification of biofilm in microtiter plates: overview of testing conditions and practical recommendations for assessment of biofilm production by staphylococci. Apmis. 2007;115(8):891–9.

19. Pompilio A, Pomponio S, Crocetta V, Gherardi G, Verginelli F, Fiscarelli E, et al. Phenotypic and genotypic characterization of Stenotrophomonas maltophilia isolates from patients with cystic fibrosis: genome diversity, biofilm formation, and virulence. BMC microbiology. 2011;11(1):1–17.

20. Helal Z, Hafez HM, Khan MI. Susceptibility of multidrug resistant Pseudomonas aeruginosa to commonly used biocides and its association with Qac efflux pump genes. Egypt J Med Microbiol. 2015;38:1–10.

21. Huang Z-m, Mi Z-h, Shi X-x, Wu L, Qin L, Wu J, et al. Disinfectant-resistant Gene of qacEΔ 1 in Gram-negative Bacteria Caused Nosocomial Infection [J]. Chinese Journal of Nosoconmiology. 2005;7.

22. Zhuo C, Zhao Q-y, Xiao S-n. The impact of spgM, rpfF, rmlA gene distribution on biofilm formation in Stenotrophomonas maltophilia. PloS one. 2014;9(10):e108409.

23. Gilbert P, Maira-Litran T, McBain AJ, Rickard AH, Whyte FW. The physiology and collective recalcitrance of microbial biofilm communities. 2002.

24. Bostanghadiri N, Ghalavand Z, Fallah F, Yadegar A, Ardebili A, Tarashi S, et al. Characterization of phenotypic and genotypic diversity of Stenotrophomonas maltophilia strains isolated from selected hospitals in Iran. Frontiers in microbiology. 2019;10:1191.

25. Azimi A, Aslanimehr M, Yaseri M, Shadkam M, Douraghi M. Distribution of smf-1, rmlA, spgM and rpfF genes among Stenotrophomonas maltophilia isolates in relation to biofilm-forming capacity. Journal of Global Antimicrobial Resistance. 2020;23:321–6.

26. Sun E, Liang G, Wang L, Wei W, Lei M, Song S, et al. Antimicrobial susceptibility of hospital acquired Stenotrophomonas maltophilia isolate biofilms. Brazilian Journal of Infectious Diseases. 2016;20:365–73.

27. Nasr AM, Mostafa MS, Arnaout HH, Elshimy AAA. The effect of exposure to sub-inhibitory concentrations of hypochlorite and quaternary ammonium compounds on antimicrobial susceptibility of Pseudomonas aeruginosa. American journal of infection control. 2018;46(7):e57–e63.

